# The relationship between neural differentiation and exploratory eye movements in healthy young and older adults

**DOI:** 10.1101/2025.01.30.635806

**Authors:** Sabina Srokova, Nehal S. Shahanawaz, Michael D. Rugg

**Affiliations:** Department of Psychology and Evelyn F. McKnight Brain Institute, University of Arizona, Tucson, AZ 85721; Center for Vital Longevity and School of Behavioral and Brain Sciences, University of Texas at Dallas, 1600 Viceroy Dr. #800, Dallas, TX 75235

## Abstract

Age-related neural dedifferentiation, characterized by lower selectivity of neural responses in the high-level sensory cortex, is thought to be a major contributor to age-related cognitive decline. However, the mechanisms that underlie dedifferentiation are poorly understood. Building on prior evidence that exploratory eye movements differ with age, we investigated the role that eye movement might play in age-related declines in neural selectivity. Healthy young and older adults of both sexes underwent fMRI with simultaneous eye-tracking as they viewed words paired with images of scenes and objects prior to a memory test. Consistent with numerous prior reports, multivoxel pattern similarity analysis (PSA) of the fMRI data revealed age-related neural dedifferentiation in scene-selective, but not object-selective cortex. Eye movements, operationalized as the number of gaze fixations made during stimulus viewing, were elevated in older adults. Analyses examining the relationship between trial-wise estimates of neural selectivity and fixation count revealed an age-invariant positive relationship in the object-selective lateral occipital complex. However, this association was found to be age-dependent in scene-selective cortex, such that there was a robust positive relationship between scene selectivity and fixation count in young, but not older adults. Lastly, scene-related selectivity in the parahippocampal place area predicted subsequent memory performance independently of age group and fixation counts at encoding. These findings suggest that age-related neural dedifferentiation in scene-selective cortex may relate to declines in the functional specialization of exploratory eye movements during scene viewing.

**Significance statement:** Healthy aging is associated with lower neural selectivity in brain regions responsive to distinct categories of perceptual stimuli. This phenomenon is most commonly observed in scene-selective cortex, although the underlying reasons remain unknown. Here, we report that trial-wise estimates of neural selectivity positively covary with the number of eye movements made during stimulus viewing. However, we also report an age-related breakdown in this relationship within scene-selective cortex, consistent with the notion that age differences in eye movements and neural selectivity reflect deficits in the extraction and binding of visual information into memory. The present findings highlight the complex interactions that occur between age, neural selectivity and eye movements, and provide novel insights into neural and behavioral mechanisms underlying cognitive aging.

## Introduction

Healthy aging is associated with neural dedifferentiation, a reduction in the neural selectivity of higher-level sensory cortex (for reviews, see Koen & Rugg, 2019; Koen et al., 2020). Prior findings indicate that lower neural selectivity is associated with poorer performance on tests of episodic memory and fluid intelligence (e.g., Park et al., 2010; Koen et al., 2019; Srokova et al., 2020; 2024) and is accompanied by higher levels of cortical tau in cognitively healthy older adults (Maass et al., 2019; Sheng et al., 2024). These findings support the proposal that age-related neural dedifferentiation significantly contributes to age-related cognitive decline. However, the mechanisms underlying dedifferentiation are poorly understood.

When operationalized at the level of stimulus categories, lower neural selectivity is consistently observed in scene-selective cortical regions (Voss et al., 2008; Carp et al. 2011; Zheng et al., 2018; Koen et al., 2019; Srokova et al., 2020, 2024), whereas null age effects are commonly reported for objects, words, or faces (Chee et al., 2006; Payer et al., 2006; Voss et al., 2008; Zheng et al., 2018; Koen et al., 2019; Srokova et al., 2020, 2024). Computational models of cognitive aging posit that lower neural selectivity reflects the broadening of neural tuning prompted by age-related alterations in the availability of cortical dopamine (Li et al., 2001, 2003) or γ-aminobutyric acid (GABA; Lalwani et al. 2019; Cassady et al., 2019, 2020; Chamberlain et al., 2021). However, because declines in neurotransmitter availability are typically considered to operate cortex wide, this assumption provides little insight into why age-related dedifferentiation should be apparent in some cortical regions but not others. As we have proposed previously (Srokova et al., 2020; 2024), a potential explanation may relate to a decline in the ability to process and bind features of relatively complex visual stimuli. Indeed, while object images depict a single item devoid of contextual information, perceiving more complex visual stimuli, such as scenes, requires integration of multiple, spatially distributed, visual features.

Eye movements support the accumulation and integration of visual information as it becomes encoded into memory (e.g., Loftus, 1972; Henderson et al., 2005; Damiano & Walther, 2019). Neuroimaging evidence suggests that more frequent gaze fixations are associated with higher stimulus-evoked fMRI BOLD signals in early and higher-level visual cortex (Henderson & Choi, 2015; Henderson et al., 2020), as well as in the hippocampus (Liu et al., 2017; 2018; 2020). In light of these findings, it is intriguing that older age is associated with marked alterations in eye movement behaviors (for review, see Ryan et al., 2019; Wynn et al., 2020). In one relevant study (Liu et al., 2018), younger and older adults underwent simultaneous fMRI with eye-tracking as they viewed face stimuli. The data revealed that older adults made more frequent eye movements (i.e., higher trial-wise fixation counts), and that the relationship between eye movements and fMRI BOLD signals in the hippocampus and fusiform face area was weaker than in younger adults. These findings suggest that age-related increases in eye movements may reflect a reduced capacity to bind visual information into a coherent memory representation.

In the present study, we employed fMRI with simultaneous eye-tracking to test the hypothesis that lower neural selectivity in older adults during memory encoding is associated with age differences in eye movements. Participants studied word-object and word-scene stimulus pairs prior to a memory test. As in our prior work (Koen et al., 2019; Srokova et al., 2020, 2024), neural selectivity was quantified using multivoxel pattern similarity analysis. We predicted that older adults would exhibit lower category-level neural selectivity for scene but not object stimuli, and this would be accompanied by an age-related increase in gaze fixation count during image viewing. Furthermore, neural selectivity was predicted to demonstrate a positive association with fixation counts in young adults, but that this association would be weaker in older adults, especially during scene viewing.

## Materials and Methods

### Participants

Twenty-six young and 32 older adults participated in the study. Two younger adults were excluded from the analyses, one due to an incidental MR finding, and another due to technical malfunction during MR acquisition. Five older adults were excluded due to poor eye-tracking calibration prior to one or more scanner runs, most commonly caused by artifacts arising from artificial intraocular lenses from a prior cataract surgery.

Participants were recruited from the University of Texas at Dallas and the surrounding Dallas metropolitan area. All participants were compensated for their time at a rate of $30/hour and up to $30 for travel. Demographic information and neuropsychological test performance for the final sample are reported in Table 1. Participants were right-handed, had normal or corrected-to-normal vision, and were fluent English speakers before the age of five. None of the participants had a history of neurological or psychiatric disease, substance abuse, diabetes, or current or recent use of prescription medication affecting the central nervous system. All participants undertook a neuropsychological test battery prior to the MRI session, and a set of inclusion and exclusion criteria were employed to minimize the likelihood of including older participants with mild cognitive impairment or early dementia (see below). All participants provided written informed consent before participation in accordance with the requirements of the Institutional Review Board of the University of Texas at Dallas.

**Table 1:** Demographic information and the outcome of the neuropsychological test battery in younger and older adults.

|  | Younger Adults | Older Adults | p-value |
| --- | --- | --- | --- |
| Age | 21.38 (3.13) | 68.56 (3.79) |  |
| Sex (male/female) | 15/9 | 7/20 |  |
| *Years of Education* | 15.19 (2.39) | 17.33 (1.92) | **0.001 \*** |
| *MMSE* | 28.71 (1.33) | 28.93 (1.00) | 0.517 |
| *CVLT Short Delay - Free* | 12.83 (2.28) | 10.22 (2.41) | **< 0.001 \*** |
| *CVLT Short Delay - Cued* | 13.13 (1.78) | 12.07 (2.13) | 0.088 |
| *CVLT Long Delay - Free* | 12.58 (2.50) | 10.78 (2.21) | **0.009 \*** |
| *CVLT Long Delay - Cued* | 12.96 (2.01) | 12.07 (2.13) | 0.134 |
| *CVLT Recognition Hits* | 15.29 (1.00) | 14.93 (1.17) | 0.235 |
| *CVLT False Alarms* | 0.63 (1.17) | 2.44 (2.29) | **< 0.001 \*** |
| *F-A-S* | 40.17 (11.62) | 44.52 (9.68) | 0.156 |
| *WMS Logical Memory I* | 30.21 (5.37) | 28.52 (4.25) | 0.223 |
| *WMS Logical Memory II* | 28.54 (5.27) | 25.93 (4.88) | 0.074 |
| *Symbol-digit Substitution Test* | 59.88 (9.08) | 50.59 (7.83) | **< 0.001 \*** |
| *Trails A (s)* | 20.70 (5.66) | 26.06 (6.34) | **0.002 \*** |
| *Trails B (s)* | 44.27 (13.01) | 61.18 (21.90) | **0.001 \*** |
| *Digit Span* | 20.33 (4.47) | 17.78 (3.81) | **0.034 \*** |
| *Category Fluency Test* | 22.75 (4.62) | 20.74 (4.64) | 0.128 |
| *TOPF* | 51.67 (9.09) | 55.74 (8.08) | 0.099 |
| *Raven's Progressive Matrices I* | 10.54 (1.50) | 10.04 (1.74) | 0.273 |
| *Visual Acuity<sup>1</sup>* | -0.02 (0.11) | 0.09 (0.12) | **0.001 \*** |

<sup>1</sup> LogMAR value

### Neuropsychological testing

Participants completed the laboratory’s standard neuropsychological test battery on a separate day prior to the MRI session. The assessment battery consists of the Mini-Mental State Examination (MMSE), the California Verbal Learning Test II (CVLT; Delis et al., 2000), Wechsler Logical Memory Tests 1 and 2 (Wechsler, 2009), the Symbol Digit Modalities Test (SDMT; Smith, 1982), the Trail Making Tests A and B (Reitan and Wolfson, 1985), the F-A-S subtest of the Neurosensory Center Comprehensive Evaluation for Aphasia (Spreen and Benton, 1977), the Forward and Backward digit span subtests of the revised Wechsler Adult Intelligence Scale (WAIS; Wechsler, 1981), the Category Fluency test (Benton, 1968), Raven’s Progressive Matrices List I (Raven et al., 2000), and the Test of Premorbid Functioning (TOPF; Wechsler, 2011). Participants also completed a visual acuity test using ETDRS charts, assessed using the logMAR metric (Ferris et al., 1982; Bailey and Lovie-Kitchin, 2013) and with corrective lenses, if prescribed. Participants were excluded prior to the fMRI session if they performed > 1.5 SD below age norms on two or more non-memory tests, if they performed > 1.5 SD below the age norm on at least one memory-based test, or if their MMSE score was < 26.

### Experimental materials

The study and test phases were completed inside the MRI scanner. Experimental stimuli were presented using PsychoPy v2021.1.3 (Peirce et al., 2019), projected onto a translucent screen (41 cm × 25 cm; 1,920 × 1,080 pixels resolution) placed at the rear of the scanner bore and viewed via a mirror mounted on the head coil (viewing distance ∼105 cm). The experimental stimuli during the study phase consisted of 128 concrete nouns, 64 colored images of objects (32 manmade and 32 natural), and 64 colored images of scenes (32 indoor and 32 outdoor). An additional 20 words, 8 scenes, and 8 objects were used as practice stimuli or as fillers at the beginning of each scanner run (one filler trial per run). The test phase included 192 concrete nouns, 128 of which were previously seen at study, and 64 of which were unstudied. Lastly, null trials (a white fixation lasting 4 sec) were randomly interspersed among the critical trials of the study (32 trials) and test phase (48 trials).

The stimulus images were presented on a gray background and were resized to fit inside a frame subtending 900 x 900 pixels. The images of objects depicted items which could be considered either “man-made” (e.g., tools, chairs, clothes) or “natural” (e.g., fruits, plants, animals). Scene exemplars reflected a variety of indoor (e.g., living room, hallway, bar) or outdoor spaces (e.g., park, field, beach). The stimulus pool described above was used to create 27 stimulus lists, 24 of which were assigned to yoked pairs of younger and older adults. Three stimulus lists were assigned to the remaining 3 older adults. For all stimulus lists, the stimuli were pseudorandomized such that participants viewed no more than three consecutive trials of the same visual category, and no more than two consecutive null trials.

### Study and test phase procedures

A schematic of the study and test tasks, including examples of experimental stimuli, are illustrated in Figure 1. Participants received instructions and completed practice study and test tasks before entering the scanner. Practice was administered on a Dell laptop computer equipped with a 17-inch display and a resolution of 1920 x 1080 pixels. Following practice, participants underwent fMRI as they completed two study-test cycles. MRI-safe glasses with appropriate lens strength were provided prior to scanning, if necessary. Each study-test cycle consisted of two study runs (∼5 minutes each) followed by two test runs (∼10 minutes each). Each study trial began with a red fixation cross presented for 500 ms, followed by a concrete noun word for 1000 ms. The word was followed by an image of a scene or an object for 4000 ms, and then a white fixation cross for an additional 1000 ms. When presented with an image of a scene, participants were asked to imagine the object denoted by the word interacting or moving around within the scene. During object trials, participants imagined the object denoted by the word and the image interacting together. Participants had a total of 5 seconds following the onset of the image to rate the vividness of the imagined scenario on a 3-point scale (“not vivid”, “somewhat vivid”, “very vivid”) using a scanner-compatible button box. The ratings were mapped onto the index, middle, and ring fingers of the right hand, respectively.

**Figure 1:**
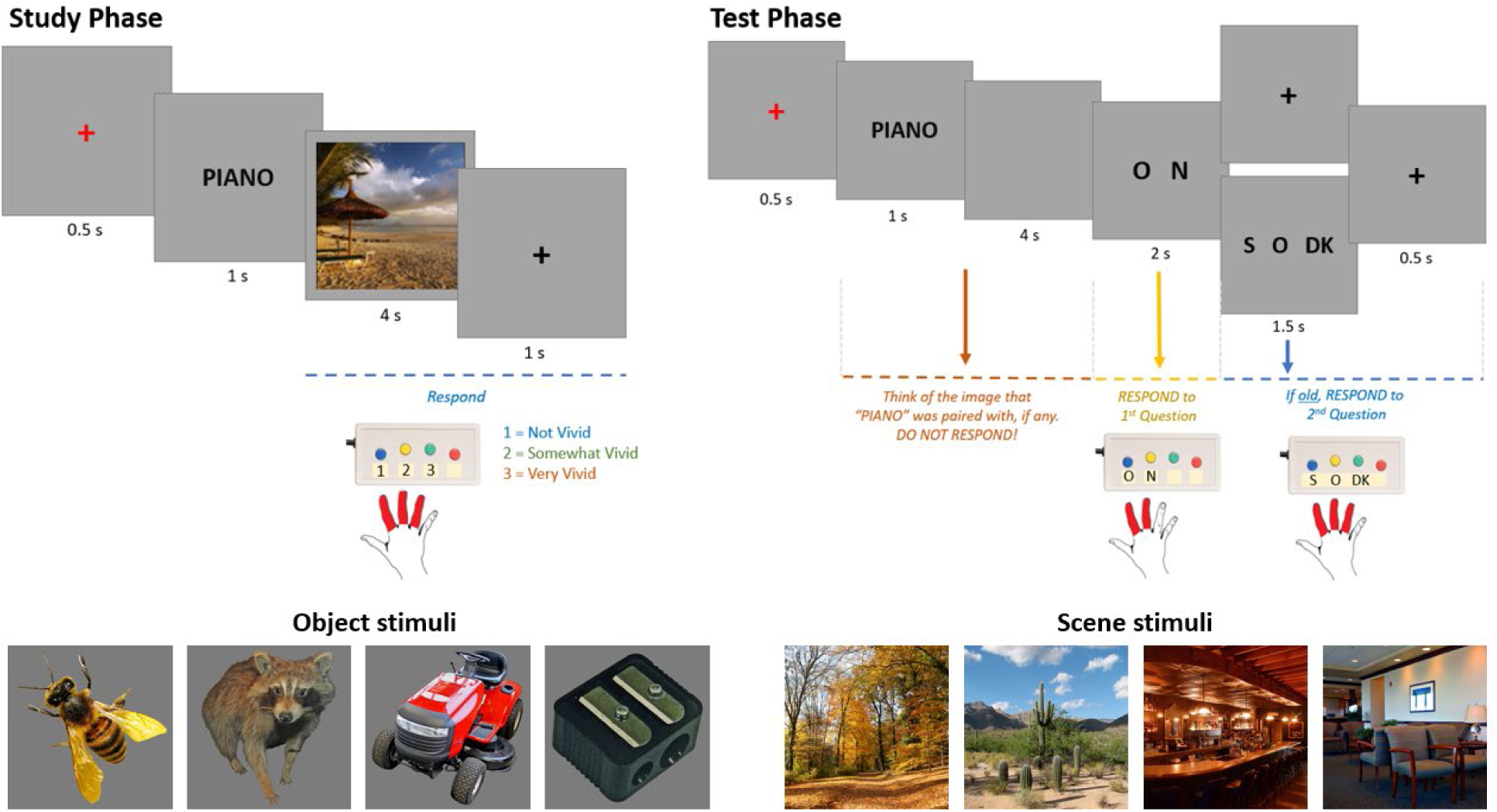
At study, participants viewed words paired with images of objects or scenes, and imagined a scenario where the object denoted by the word was interacting with the image. At test, participants viewed old and new words. If old, participants were asked to think of the image that had been paired with the word. After the imagery period, participants indicated whether the word had been old or new, and if old, whether it had been paired with a scene or an object. Four example object and scene stimulus images are illustrated below the task schematics.

Each trial of the test phase began with a red fixation cross presented for 500 ms, followed by a word for 1000 ms. Next, participants viewed a blank screen for 4000 ms, during which they were asked to imagine the image that had been paired with the test word, if any. Following the blank screen period, participants indicated whether they remembered seeing the word by making an “Old” or “New” response. For each word endorsed “Old”, participants went on to make a source memory judgement, indicating whether the word had been paired with a scene or an object. A third “Don’t Know” option was included to discourage guessing. The Old/New judgements were made within 2 seconds using the index and middle fingers of the right hand. The Scene/Object/Don’t know judgements were also made within 2 seconds, this time using the index, middle, and ring fingers of the right hand. In both cases, the ordering of the response options was counterbalanced across participants, while ensuring that the Don’t know response was never assigned to the middle finger. Following the response period, each test phase trials concluded with a white fixation cross presented for 500 ms.

### Eye movement recording

Eye movement data were acquired simultaneously with fMRI using an MRI-compatible eye-tracker (Eyelink 1000 Plus; SR Research). Monocular data was acquired from the right eye at a sampling rate of 1000 Hz. Calibration and manual validation were performed using Eyelink’s 9-point calibration procedure at the beginning of each scanner run, motivated by evidence that, due to restricted head movement in the scanner, the drift in eye-tracking accuracy is minimal when calibration is performed prior to each run (Peitek et al., 2018). The calibration/validation procedure was iterated prior to each scanner run until the accuracy of validation at each gaze point was acceptable as per Eyelink’s default threshold. As noted above, five older adults were excluded from the present study as we were unable to accurately calibrate and validate their gaze maps. Eyelink’s default event parser was used to classify eye movements into blinks, saccades, and fixations. A velocity threshold of 30°/s and an acceleration threshold of 8000°/s^2^ were used to identify saccades. Samples with missing pupil size data were labeled as blinks. The remaining data were classified as fixations.

### Eye-tracking analyses

Eye movement data was recorded in an EDF format which was subsequently imported into MATLAB using the edfmex.m function (Kovach, 2007). Custom MATLAB scripts were used to calculate the number of gaze fixations during the 4 seconds when the stimulus image remained on the screen. We excluded fixations that fell outside of the coordinates spanning the 900x900 pixel stimulus boundaries (fixations that fell 1-degree visual angle outside of the boundaries were kept to account for potential error in gaze position). Additionally, to ensure that the analyses were not confounded by potential age-related decrease in gaze stability, we removed all fixations that were shorter than 50ms. We also performed an exploratory eye movement dispersion analysis, where dispersion was computed as the average Euclidean distance between the coordinates of each fixation occurring during a given trial.

### Exploratory pupillometry analyses

Although the primary focus on the present paper was on eye movement behaviors (fixation counts), we took advantage of the availability of the pupillometry data to form an exploratory analysis examining any potential age differences in pupil size at encoding. Pupil data was analyzed with custom MATLAB code. Firstly, the data were de-blinked such that any samples belonging to blink events were removed (set to NaN), along with any samples occurring within 80ms before/after a blink. The median absolute deviation method (Kret & Sjak-Shie, 2019) was employed to filter dilation speed outliers (i.e., changes in pupil size that are improbable given the typical pupil dilation/constriction speed). Next, the data was smoothed with a 100-point moving average window and missing data was estimated using linear interpolation, with the restriction that data were not interpolated over gaps larger than 1000 ms. Lastly, the data were baseline-corrected by subtracting the raw pupil size acquired during the initial 500 ms of a given trial. Trials containing < 33% of samples prior to interpolation, or < 50% during baseline, were excluded.

### MRI data acquisition and preprocessing

Functional and structural MRI data were acquired at the Sammons BrainHealth Imaging Center at the University of Texas at Dallas with a Siemens Prisma 3T scanner. The scanner was equipped with a 32-channel head coil. A whole-brain anatomical scan was acquired with a T1-weighted 3D MPRAGE pulse sequence (FOV = 256 × 256 mm; voxel size = 1 × 1 × 1 mm; 160 slices; sagittal acquisition). Functional data were acquired with a T2*-weighted BOLD echoplanar imaging (EPI) sequence with a multiband factor of 3 (flip angle = 70° FOV = 220 × 220 mm; voxel size = 2 × 2 × 2 mm; TR = 1.52 ms; TE = 30 ms; 66 slices). A dual-echo fieldmap sequence which matched the 3D characteristics of the EPI sequence was acquired at TEs of 4.92 and 7.38 ms immediately after the last run of the study phase, resulting in two magnitude images (one per echo), and a pre-subtracted phase image (the difference between the phases acquired at each echo).

The MRI data were preprocessed using Statistical Parametric Mapping (SPM12, Wellcome Department of Cognitive Neurology) and custom MATLAB code (MathWorks). The functional data were preprocessed in six steps. First, we employed the FieldMap toolbox in SPM to calculate voxel displacement maps prior to the fieldmap correction. These maps were calculated using the magnitude and phase difference images acquired in the aforementioned dual-echo fieldmap sequence. Second, SPM’s realign and unwarp procedure was applied, operating in two steps: spatial realignment of the time series registered to the mean EPI image and a dynamic correction of the deformation field using the voxel displacement maps. Third, the functional images were reoriented along the anterior and posterior commissures, then spatially normalized to SPM’s EPI template, and then renormalized to an age-unbiased sample-specific EPI template according to procedures standardly employed in our laboratory (de Chastelaine et al., 2016). Lastly, the functional data were smoothed with a 5 mm FWHM Gaussian kernel.

### Whole-brain univariate and parametric modulation analyses

The functional data were analyzed with a two-stage univariate GLM approach. At the first stage, separate GLMs were implemented for the study data of each participant. The study trials were binned into two events of interest (scene and object trials), and the neural activity elicited by the trials was modeled with a 4 second boxcar extending throughout the presentation of the stimulus image. The boxcar regressors were convolved with SPM’s canonical hemodynamic response function (HRF). Filler trials, trials which did not receive a vividness response, or which received a response earlier than 500ms post-image-onset were included as events of no interest. Additional covariates of no interest comprised 6 motion regressors reflecting rigid-body translation and rotation, spike covariates regressing out volumes with transient displacement of > 1 mm or > 1 ° in any direction, and the mean signal of each scanner run. The parameter estimates from the first-level GLM were carried over to a 2 (age group) x 2 (image category) mixed factorial ANOVA. The outcomes of the group-level analyses were height-thresholded at p < 0.05 (FWE corrected) with a voxel extent threshold of k = 50.

Parametric modulation analysis was employed to identify the brain regions where number of gaze fixations covaried with fMRI BOLD activity during image viewing. At the first stage, separate GLMs were performed for each individual subject in an approach similar to the whole-brain univariate analyses. Two events of interest (scenes, objects) and trials of no interest were entered into the model as 4 second boxcars and convolved with canonical HRF. For each event of interest, mean-centered vectors of the trial-wise number of fixations were entered as the parametric modulator. Regressors of no interest comprised 6 motion regressors, motion spike covariates, and the mean signal of each scanner run. The parameter estimates from the individual subject-level GLMs were entered into a group-level ANOVA from which three t-contrasts were derived: the main effect of gaze fixations for scenes, objects, and both image categories. Following Liu et al. (2018), we thresholded these contrasts at p < 0.005 (unc.) with a voxel extent threshold of k = 20. Age differences in modulation effects were examined in the two a priori regions-of-interest (ROIs): the parahippocampal place area (PPA) and the lateral occipital complex (LOC; *see below for ROI definition*).

### Multivoxel pattern similarity analyses

Neural differentiation for scenes and objects was examined with pattern similarity analysis (PSA). For the purposes of this analysis, the data from the study phase were subjected to a “least-squares-all” GLM (Rissman et al., 2004; Kriegeskorte et al., 2008; Mumford et al., 2014). Each trial was modeled by a separate 4 second boxcar regressor tracking the image presentation. Six motion regressors reflecting translational and rotational displacement, along with the mean signal of each session were included as covariates of no interest. Category-level PSA of the study data was conducted using approaches analogous to those employed in prior work from our laboratory (Koen et al., 2019; Srokova et al., 2020; Hill et al., 2021; Srokova et al., 2024). To ensure that similarity metrics were not confounded by within-run autocorrelation effects (Mumford et al., 2014), all correlations were computed between trials belonging to different scanner runs. PSA was operationalized for each image category (scenes/objects) by computing the differences between within-category and between-category similarity metrics. Within-category similarity was computed as the average voxel-wise Fisher’s *z*-transformed correlation between a given trial and all trials of the same image category. Between-category similarity was calculated in an analogous fashion—as the average Fisher’s *z*-transformed correlation between a given trial and all trials of the alternate image category.

### Region of interest and whole-brain parcellation analyses

Multivoxel PSAs were performed at the ROI and whole-brain levels. We followed our previously published approach to define two a priori ROIs – the scene-selective parahippocampal place area (PPA) and the object-selective lateral occipital complex (LOC; Koen et al., 2019; Srokova et al., 2024). To ensure that the ROIs were derived independently from the to-be-analyzed data, and that they were unbiased with respect to age group, the ROIs were generated using an independent dataset where younger and older adults viewed scene and object stimuli during encoding (see Srokova et al., 2024 for the dataset). The data in the independent study were analyzed with a two-stage univariate GLM approach. At the individual subject level, events of interest associated with object and scene stimuli were modeled with a 2 s boxcar function encompassing the image presentation and convolved with SPM’s canonical HRF. The individual models covaried out the six motion regressors reflecting rigid-body translation and rotation, as well as spike covariates regressing out volumes with displacement greater than 1mm or 1° in any direction, and the mean signal of each run. At the second level, the PPA was defined with a directional novel scene > novel object t-contrast collapsed across both age groups, and thresholded at p < 0.05 (FWE-corrected). The scene-selective contrast was inclusively masked with the anatomical labels of the parahippocampal and fusiform gyri provided in SPM’s Neuromorphometrics atlas, and restricted with a 15 mm radius sphere centered around the peak voxel in order to separate the PPA from scene-selective effects in the subiculum. An analogous approach was used to define the LOC, using the object > scene contrast masked with the labels of the middle and inferior occipital gyrus. The final sizes of the ROIs was: right PPA = 254 voxels, left PPA = 279 voxels, right LOC = 350 voxels, left LOC = 296 voxels. The inclusion of hemisphere as a factor did not alter the outcome of any of the analyses reported in the present manuscript. Therefore, the analyses were performed after averaging across the left and right hemispheres to generate a single bilateral ROI.

The ROI-based PSA was complemented with a whole-brain analysis. Although age differences in neural selectivity are commonly examined with an ROI-based approach, PSA effects are unlikely to be restricted only to those regions that exhibit univariate selectivity. Therefore, we conducted a whole-brain PSA using the MVPA approaches described above on cortical parcels defined by the Schaefer atlas (Schaefer et al. 2018). The analysis was performed using the parameter estimates extracted from each parcel. The atlas was masked with a group-level mask generated from the present dataset, and parcels that contained less than 10 voxels were excluded from analyses. This resulted in a final number of 996 parcels. In the first pass of the analyses, we identified regions that exhibited significant similarity effects (i.e., significantly different from zero) separately in both age groups using an FDR-adjusted significance threshold of q < 0.05. Age differences in selectivity were examined only in the regions exhibiting reliable similarity effects in both age groups.

### Linear mixed effects analyses

The relationship between PSA measures of neural differentiation and trial-wise fixation counts was examined using a series of linear mixed effects (LME) analyses performed at both the ROI and whole-brain levels. In the present context, LMEs are ideal for studying how trial-wise fluctuations in neural selectivity and eye movement metrics covary by explicitly accounting for within-subject variability and individual differences across trials. By modeling subject-specific intercepts and/or slopes, LMEs capture dynamic, trial-by-trial changes within each subject, providing more precise and reliable estimates of fixed effects without averaging out important fluctuations. For analyses of the scene trials, tests were restricted to the PPA ROI and those Schaefer parcels that exhibited main effects of scene similarity. Similarly, object analyses were restricted to the LOC and all object-selective parcels. In each selected parcel, we first tested for an interaction between age and fixation count in the prediction of neural selectivity by employing the following syntax: *PSA similarity ∼ age group * fixation count + (1 | subject).* We report random intercept-only models here given that not all Schaefer parcels converged when random slopes were included. However, we note that the inclusion of random slopes in the PPA and LOC ROIs did not alter the statistical outcomes of any of the analyses reported below. In cases where the interaction was not significant (i.e., the relationship between neural selectivity and fixation count did not differ by age group), the LME was re-run using the syntax of *PSA similarity* ∼ *age group + fixation count + (1 | subject)*. This model tested whether fixation count explained unique sources of variance in PSA similarity across both age groups. In cases where the interaction was significant (i.e., the relationship differed between young and older adults), *PSA similarity ∼ fixation count + (1 | subject)* models were run separately for younger and older adults. Trials where the total number of fixations equaled zero were removed from all mixed effects analyses.

### Relationship with subsequent memory performance

Binomial generalized LMEs were employed to examine the relationship between subsequent memory success, age group, fixation count, and PSA similarity in the PPA (scene trials) and LOC (object trials). Memory success was binarized into correct (source correct) and incorrect trials (i.e., source incorrect, source don’t know, and item miss trials) prior to the analysis. Given that no significant interactions with age group were identified in any of the generalized LMEs reported below (ps > 0.163), and to avoid overfitting and convergence issues, the syntax used to generate all statistical outcomes was: *memory bin ∼ age group + PSA similarity + fixation count + (1 | subject)*.

## Results

### Neuropsychological test performance

Neuropsychological test performance is illustrated in Table 1. Young adults outperformed older adults on the following measures: CVLT Free recall (short delay and long delay), CVLT false alarms, SDMT, Trails A and B, digit span, and visual acuity. There were no measures where older adults performed better than their younger counterparts.

### Vividness ratings, but not reaction times, during encoding are moderated by age group

Two 2 (age group) x 2 (category) mixed-effects ANOVAs were performed to examine potential categorical and age differences in vividness ratings and reaction times (RTs) during the study phase. Vividness ratings were missing from one older adult participant due to technical malfunction. The ANOVA on these ratings revealed a significant main effect of age group (F(1,48) = 33.556, p < 0.001, partial *η*^2^ = 0.411; higher vividness ratings in younger adults), a main effect of category (F(1,48) = 18.257, p < 0.001, partial *η*^2^ = 0.276; higher vividness ratings for scenes), and an age group x category interaction (F(1,48) = 4.465, p = 0.040, partial *η*^2^ = 0.085). Follow-up t-tests indicated that age differences in vividness ratings were observed for both stimulus categories, (Objects: younger mean [SD] = 2.377 [0.400]; older mean [SD] = 1.785 [0.324]; t(44.36) = 5.722, p < 0.001, d = 1.633); Scenes: younger mean [SD] = 2.449 [0.305]; older mean [SD] = 1.999 [0.322]; t(47.96) = 5.079, p < 0.001, d = 1.435), but with greater age differences evident for objects than scenes (mean difference = 0.592 and 0.45).

The analogous ANOVA of RTs revealed a null effect of age group (F(1,48) = 0.026, p = 0.872, partial *η*^2^ = 0.001), a significant effect of category (F(1,48) = 4.896, p = 0.032, partial *η*^2^ = 0.093), reflective of longer RTs for scenes (mean [SD] = 2.923s [0.911]) than objects (mean [SD] = 2.877 [0.916]), and a null age by category interaction (F(1,48) = 0.978, p = 0.328, partial *η*^2^ = 0.020). Together, these behavioral results indicate that while older adults rated the imagined scenarios as less vivid, they were equally fast at making their vividness responses.

### Item recognition and source memory performance are lower in older than younger adults

Item recognition and source memory performance are summarized in Figure 2A. Item recognition was operationalized as the difference between the hit rate (the proportion of items correctly endorsed “old”) and the false alarm rate (the proportion of new items incorrectly endorsed “old”). A Welch’s two sample t-test revealed that recognition was higher in the younger (Mean [SD] = 0.719 [0.162]) than the older adult group (Mean [SD] = 0.588 [0.182]; t(48.99) = 2.716, p = 0.009, d = 0.756). Source memory performance (pSR) was estimated with a single high-threshold model (Snodgrass and Corwin, 1988) using the formula: pSR = [pSource Correct – 0.5 * (1 – pDon’t Know)]/[1 – 0.5 * (1 – pDon’t Know)], where “pSource Correct” and “pDon’t Know” refer to the proportion of correctly recognized old trials receiving an accurate source memory judgment or a “Don’t Know” response, respectively. A Welch’s two sample t-test revealed greater pSR scores in younger (Mean [SD] = 0.608 [0.215]) than older adults (Mean [SD] = 0.331 [0.230]; t(48.89) = 4.431, p < 0.001, d = 1.238). Thus, both item and source memory performance were lower in older relative to younger adults, albeit with a markedly greater effect size for source memory.

**Figure 2:**
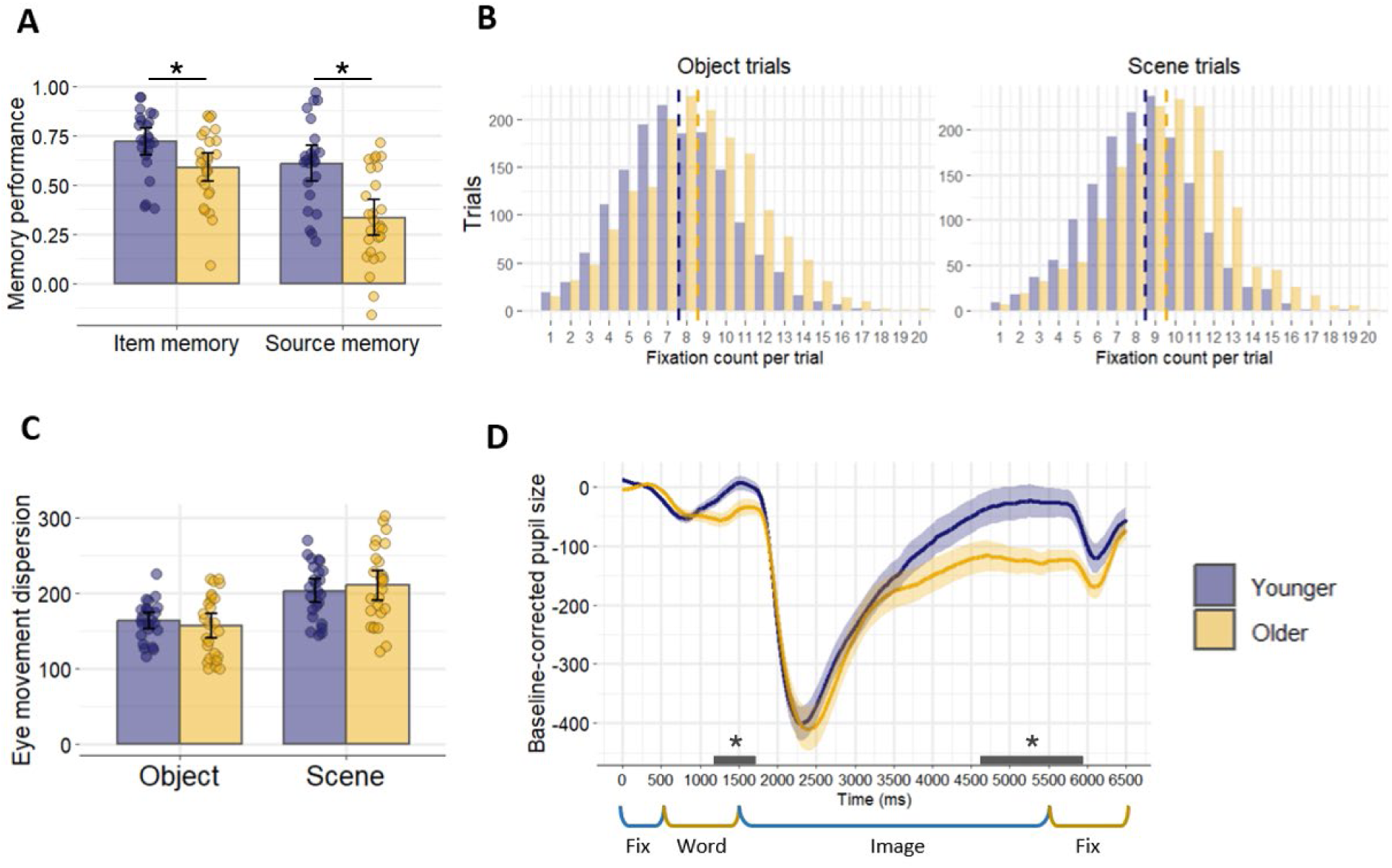
**A**: Item recognition and source memory performance in younger and older adults. Error bars reflect 95% confidence intervals. **B:** The distributions of the number of fixations performed in a given trial. Dashed lines reflect the mean fixation count for each age group, revealing that, on average, older adults performed more fixations across object and scene trials. **C:** Eye movement dispersion in younger and older adults operationalized in terms of the average Euclidean distance (pixels) between all fixations in a given trial. Error bars reflect 95% confidence intervals. **D:** Average pupil size (baseline-corrected) across the entire encoding trial. Horizontal lines represent time points (minimum duration 500 ms) during which pupil size significantly differed between younger and older adults. Shaded areas in the timeseries plot represent the standard error of the mean.

### Older age is associated with more frequent gaze fixations, but the distances between fixations do not differ by age group

For each trial, we calculated the number of gaze fixations that were made during the 4 seconds that the stimulus image was on the screen. The distributions of fixation counts are illustrated in Figure 2B. Across-trial fixation counts were entered into a 2 (age group) x 2 (category) ANOVA, which revealed a significant main effect of age group (F(1,49) = 4.529, p = 0.038, partial *η*^2^ = 0.085; older adults made more fixations), and a main effect of category (F(1,49) = 113.909, p < 0.001, partial *η*^2^ = 0.699; more fixations were made during scene viewing). The 2-way interaction was not significant (F(1,49) = 0.589, p < 0.446, partial *η*^2^ = 0.012), indicating that the older group made more fixations than their younger counterparts across both stimulus categories.

As described above (see Methods), the exploratory fixation dispersion metric was computed as the average Euclidean distance between the pixel coordinates of each fixation that occurred while the stimulus remained on the screen (Figure 2C). A 2 (age group) x 2 (category) ANOVA resulted in a null effect of age group (F(1,49) = 0.001, p = 0.984, partial *η*^2^ = 0.001), a significant main effect of category (F(1,49) = 252.530, p < 0.001, partial *η*^2^ = 0.837; higher dispersion during scene viewing), and a significant 2-way interaction (F(1,49) = 6.114, p = 0.017, partial *η*^2^ = 0.111) The follow-up two sample t-tests comparing younger and older adults separately during scene and object viewing were not significant (scenes: t(47.44) = -0.577, p = 0.567, d = -0.159; object: t(44.57) = 0.786, p = 0.436, d = 0.215), suggesting that the interaction arose from numerically (non-significantly) higher dispersion in older adults during scene viewing, accompanied by lower dispersion during object viewing. In sum, the dispersion findings indicate that although older adults moved their eyes around more frequently (higher fixation counts), the mean dispersion of eye movements did not significantly differ across the two age groups. Since older adults made more fixations overall, the fact that distances between fixations were equal across age group may suggest that additional eye movements were used to sample a broader area of the stimulus, potentially compensating for age-related decline in the functional field of view (Sekuler et al., 2000). Additionally, these results confirm that our analyses were unlikely to have been confounded by less stable fixations in older adults.

### Younger adults exhibit greater pupil dilation during stimulus viewing

Pupil timeseries data were collapsed across image categories given that the scenes and objects were not matched in luminance, making any potential category effects would be uninterpretable. (We note that similar patterns of age differences arise if the two categories are analyzed separately). The resulting across-trial pupil size for younger and older adults is illustrated in Figure 2D (Significant differences between age groups were defined as continuous time windows exhibiting differences at p < 0.05 (Welch’s two sample t-test) for at least 500ms. Pupil size was significantly lower in older than younger adults from 1211ms to 1728ms post-trial onset, coinciding with the presentation of the stimulus word. Image presentation evoked a robust pupil constriction that did not differ between age groups, indicating that the pupillary responses to changes in luminance were similar between young and older adults. While the image remained on the screen, gradual dilation of the pupil was observed. Throughout 4647ms to 5973ms post-trial onset (coinciding with image presentation), older adults’ pupil size was significantly smaller relative to the younger adults.

### Fixation count positively covaries with fMRI BOLD signals in canonical scene-selective and object-selective cortical regions

The results of the whole-brain univariate encoding and parametric modulation contrasts are described in Table 2 and illustrated in Figures 3A-B. Turning first to scene-selective effects (scene > object, collapsed across age groups), a large contiguous cluster encompassed regions of the parahippocampal and fusiform gyri, anterior hippocampus, lingual gyrus, retrosplenial complex, occipital pole, superior occipital gyrus, and the transverse occipital sulcus (bilaterally in all cases). Additional clusters were identified in bilateral precuneus and the posterior cingulate cortex, left lateral geniculate nucleus, septal nuclei, and two bilateral clusters in the inferior posterior cerebellum. Object-selective effects were evident in inferior and middle occipital areas, extending into the posterior parts of the inferior temporal lobes. Object-selective effects were also observed in bilateral inferior parietal regions, extending into the intraparietal sulci and the supramarginal gyri. Two additional clusters were present in the right inferior posterior cerebellum, and the left inferior and middle frontal gyrus.

**Figure 3:**
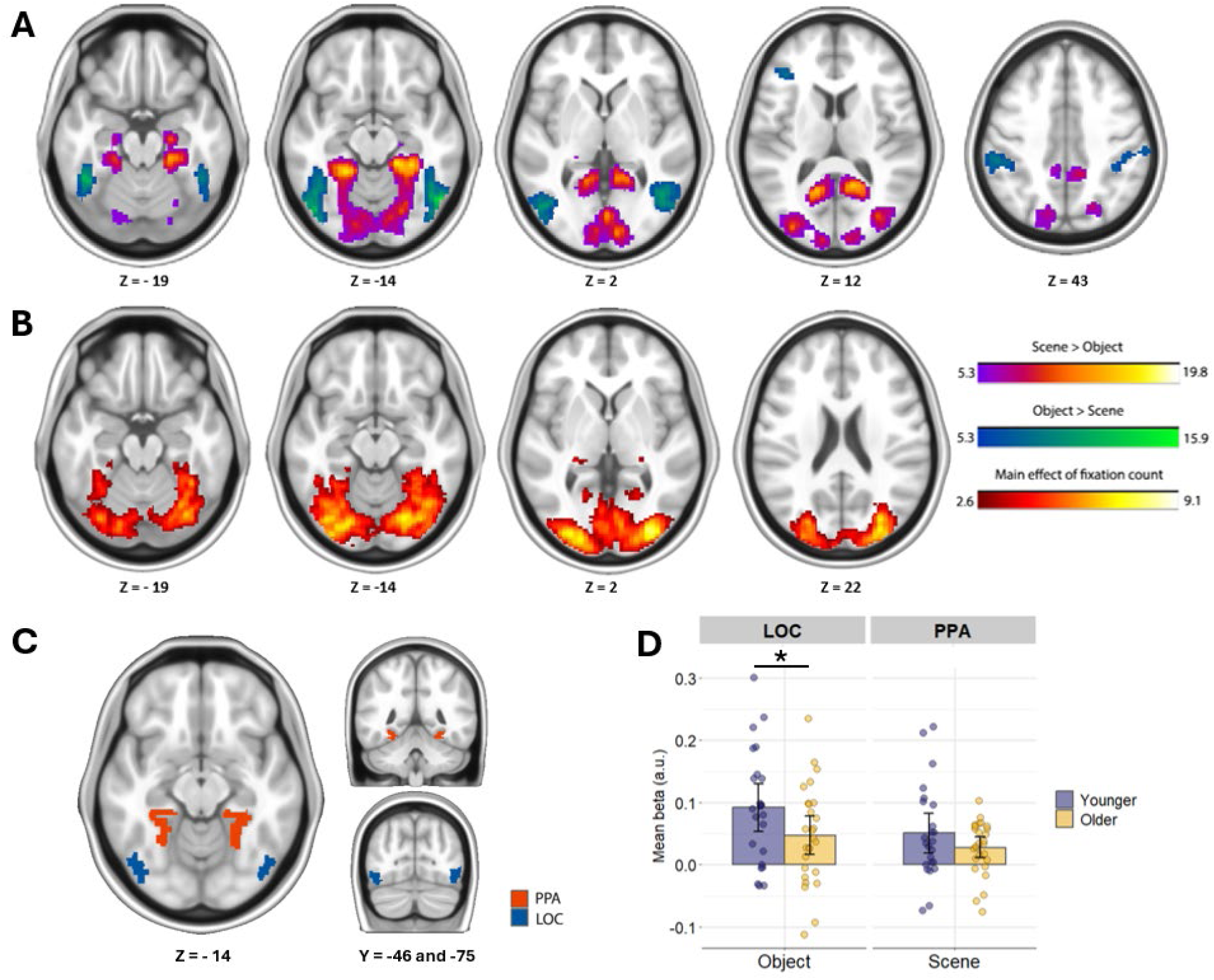
**A**: Outcomes of the across-age scene > object and object > scene t-contrasts. **B:** Positive main effects of fixation count illustrating the regions where higher fMRI BOLD covaried with more frequent eye movements. **C:** Two a priori ROIs (PPA and the LOC). **D:** Parameter estimates averaged within the PPA and LOC derived from the parametric modulation analysis. Error bars represent 95% confidence intervals. FMRI results and the ROIs are illustrated on a T1-weighted ICBM 152 MNI brain.

**Table 2:**
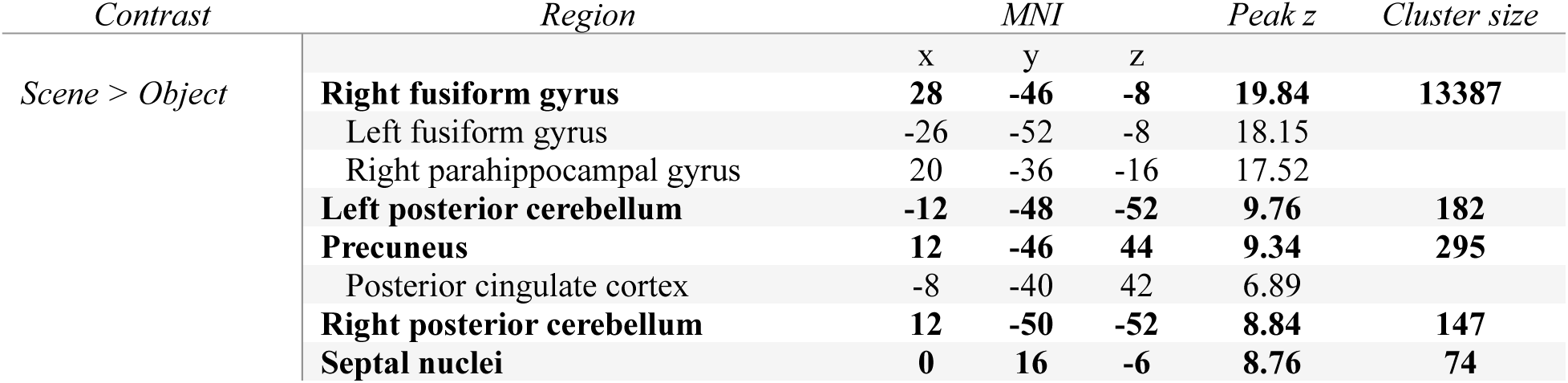
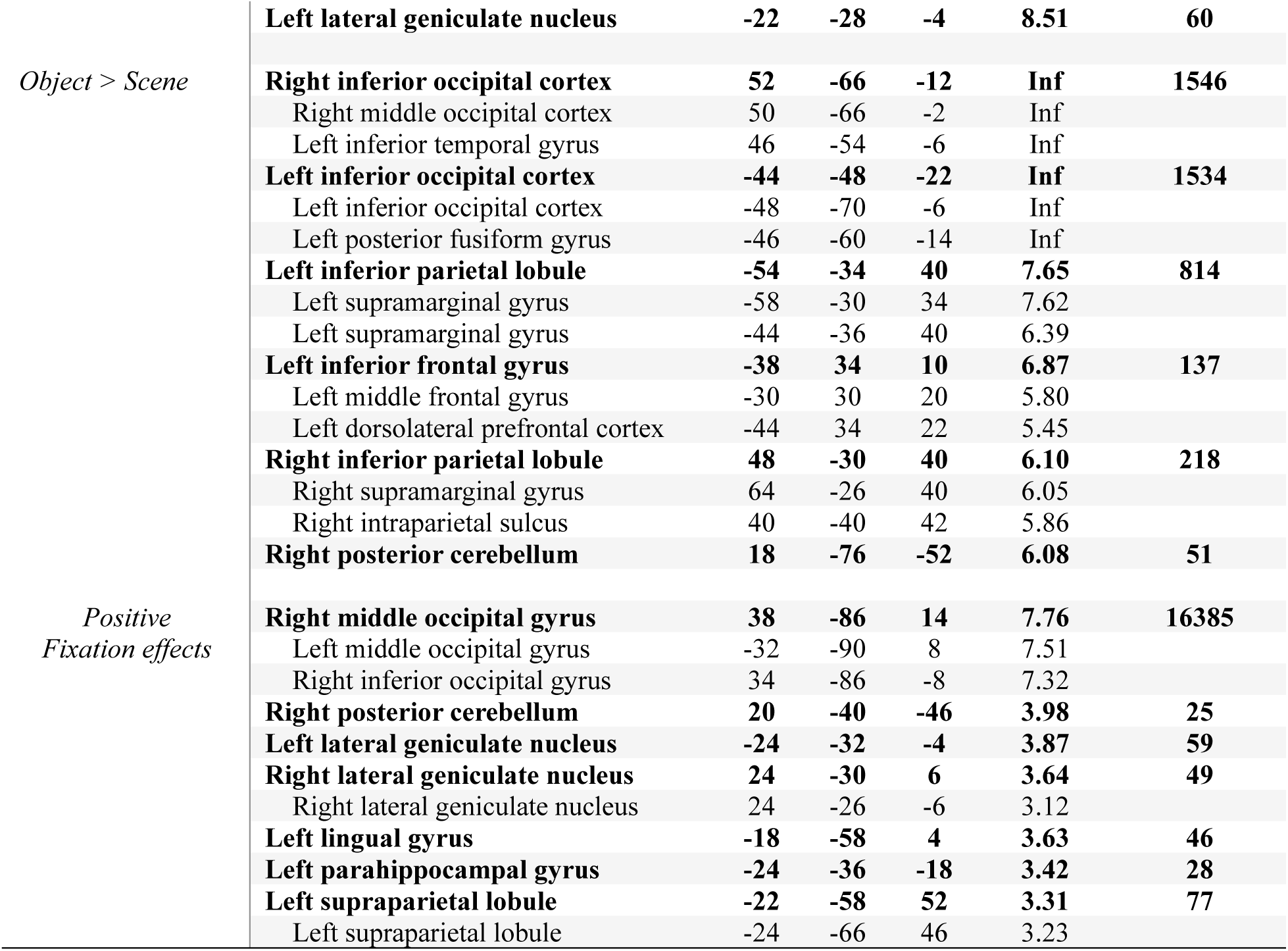
Loci of univariate scene-selective and object-selective effects, and the positive main effect of fixation count.

Turning next to the parametric modulation analyses, positive fixation effects during the scene and object trials revealed numerous highly overlapping clusters, suggesting that increased frequency of eye movements leads to an increase of fMRI BOLD activity in similar regions, regardless of the image category. Therefore, Figure 3B illustrates positive fixation effects collapsed across the two category regressors. Fixation count during stimulus viewing positively covaried with fMRI BOLD activity in a large cluster spanning the occipital pole, inferior and middle occipital areas, the fusiform and posterior parahippocampal gyri (bilaterally in all cases). Additional effects were evident in the bilateral lingual gyri, the lateral geniculate nuclei, posterior parietal areas extending into the intraparietal sulci, and the right posterior cerebellum. Importantly, greater fixation count was positively associated with greater fMRI BOLD signals in many of the scene- and object-selective regions identified in the whole-brain univariate analyses identified above. Therefore, we followed up the whole-brain modulation analyses by extracting parameter estimates from the two a priori ROIs (Figure 3C). Parameter estimates from the LOC were extracted using the regressor which corresponded to the parametric modulator of fixation counts during object viewing, whereas the parameter estimates from the PPA reflected modulation effects during scene viewing. A (2) age group x 2 (ROI) ANOVA revealed a main effect of age group (F(1,49) = 4.654, p = 0.036, partial *η*^2^ = 0.087) and a main effect of ROI (F(1,49) = 5.086, p = 0.029, partial *η*^2^ = 0.094). The interaction was not significant (F(1,49) = 0.646, p = 0.426, partial *η*^2^ = 0.013; Figure 3D). These results indicate that the modulatory effects of eye movements on fMRI BOLD signals were greater in the LOC than the PPA, and in younger relative to older adults.

### Age differences in neural selectivity are restricted to scene-selective cortex

Figure 4A depicts the parcels exhibiting significant scene- and object-related selectivity in both age groups. Scene effects were evident across the parahippocampal and fusiform gyri (PPA), the inferior and superior occipital cortex (commonly referred to as the ‘occipital place area’ - OPA), lingual gyrus, and the retrosplenial complex (often referred to as the ‘medial place area’ – MPA). Object effects were present predominantly in lateral occipital regions, while negative object effects (similarity indices significantly below zero, i.e., trials where images of objects elicited neural patterns that were more similar to scenes relative to other objects) were observed in the occipital cortex. Age differences were tested in the parcels identified in Figure 4A, and the outcomes of these analyses are illustrated in Figure 4B. For scene trials, younger adults demonstrated higher scene selectivity in several of the canonical scene-responsive regions in the occipital cortex, parahippocampal and fusiform gyri, the right MPA, and the right OPA. For object trials, younger adults exhibited higher selectivity in a single parcel within the most anterior portion of the LOC (illustrated in orange). Age differences for objects were also identified in two additional parcels in the occipital and posterior parahippocampal areas. However, these effects arose from regions exhibiting significant negative object effects and thus are not discussed further. In summary, age-related neural dedifferentiation was most reliably observed during scene processing in scene-selective cortical regions.

**Figure 4:**
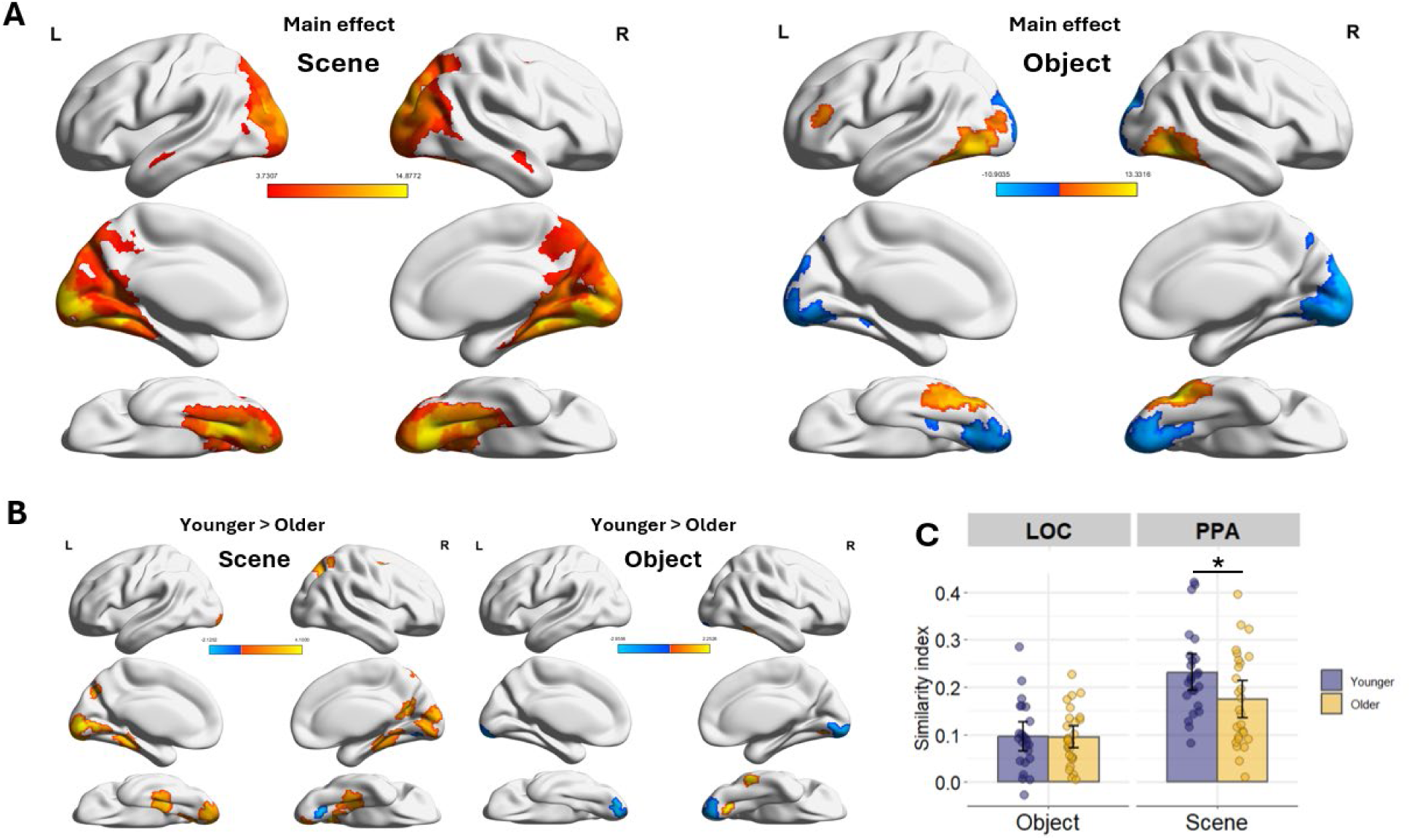
**A:** Significant scene and object PSA effects identified across young and older adults. The t-values reflect tests performed across the full sample. **B:** Regions exhibiting reliable age differences in neural similarity for scenes and objects (red, greater similarity for younger adults; blue, greater similarity for older adults**). C:** Similarity indices in the LOC and PPA ROIs during object and scene viewing, respectively, demonstrating reduced neural differentiation for scenes in the PPA. Error bars represent 95% confidence intervals.

The outcomes of the ROI analyses are depicted in Figure 4C. Similarity indices in the LOC (for objects) and PPA (for scenes) were reliably different from zero in both younger and older adults (ps < 0.001). A 2 (age group) x 2 (ROI) mixed effects ANOVA revealed a significant effect of ROI (F(1,49) = 51.323, p < 0.001, partial *η*^2^ = 0.512), but no effect of age group (F(1,49) = 2.832, p = 0.099, partial *η*^2^ = 0.055) and only a marginal trend towards an interaction (F(1,49) = 3.374, p = 0.072, partial *η*^2^ = 0.064). Given that we have demonstrated in three separate fMRI experiments that age differences in neural differentiation are more pronounced in scene-selective than in face-or object-selective regions (Koen et al., 2019; Srokova et al., 2020; 2024), we had an *a priori* justification to test for age differences separately in the LOC and the PPA. Consistent with prior findings, Welch t-tests identified lower neural differentiation in the older age group in the PPA (t(48.94) = 2.130, p = 0.038, d = 0.609), but not in the LOC (t(44.701) = 0.078, p = 0.937, d = 0.023). In summary, the whole-brain and ROI analyses provide convergent evidence that age-related neural dedifferentiation is most evident during scene viewing.

### Greater trial-wise estimates of neural selectivity are associated with higher fixation counts, and this relationship is moderated by age in the scene-selective cortex

The LME analyses described below examined whether trial-wise fluctuations of neural selectivity covaried with eye movements (fixation count) at the whole-brain or ROI levels. As noted in the Methods section, whole-brain analyses were restricted to those Schaefer parcels that exhibited significant main effects of scene or object similarity in both age groups. In other words, when examining the relationship between eye movements and neural selectivity during scene viewing, we focused the analyses only on those regions that exhibited scene-selectivity in the PSA analyses described above (and analogously for object trials; see Figure 4A).

Turning first to scene trials, Figure 5A illustrates those scene-selective parcels where fixation count positively covaried with similarity indices across age groups using the formula *PSA similarity ∼ age group + fixation count + (1 | subject)*. As is evident in the figure, greater fixation count was associated with higher scene similarity indices in most of scene-selective cortex. LME models which tested for age group by fixation count interactions in predicting PSA similarity (*PSA similarity ∼ age group * fixation count + (1 | subject)*) identified several parcels in the parahippocampal, retrosplenial, and superior occipital cortices (spatially aligning with the PPA, MPA, and OPA; Figure 5B). Significant interaction effects suggest that the relationship between eye movements and scene selectivity differed between young and older adults in these parcels. Therefore, we performed follow-up analyses separately for the two age groups, revealing that the interaction effects were driven by robust significant positive relationships in younger adults in all of the parcels identified by the interaction effect, but non-significant associations in older adults (except for the two parcels outlined in Figure 5B which, although exhibiting small effect sizes, were also significant in older adults: ps = 0.045 and 0.047). In conclusion, trial-wise fixation count was positively associated with scene similarity indices in most of scene-selective cortex. However, the findings indicate that this relationship was stronger in younger than older adults across many canonical scene-selective regions, including the PPA, MPA, and OPA.

**Figure 5:**
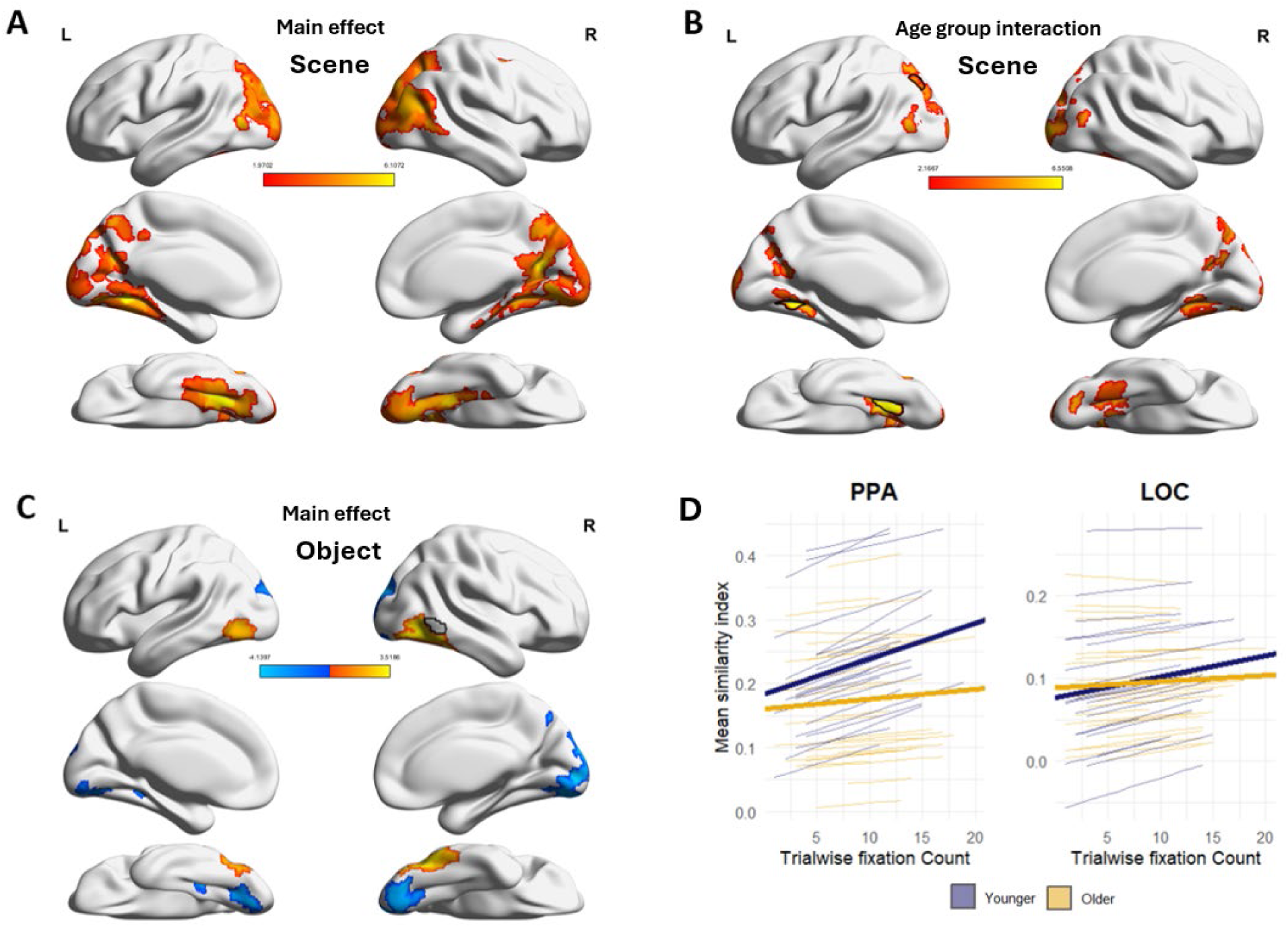
**A:** Scene-selective regions where trial-wise PSA similarity indices positively covaried with the with fixation counts. The heatmap indicates the t-values associated with the predictor of fixation count. **B:** Scene-selective regions where fixation count interacted with age group in predicting trial-wise PSA similarity indices. The parcels and t-values illustrated here reflect regions that exhibited a significant positive relationship in younger adults. In older adults, the association between eye movements and similarity indices was not significant in all but two of these parcels (outlined in black; see text for statistics). **C:** Object-selective regions where trial-wise PSA similarity indices covaried with fixation count. One parcel (outlined in black) exhibited a significant age group by fixation count interaction. **D:** Outcomes of LME analyses in PPA and LOC ROIs. PPA exhibited a significant fixation count by age group interaction, reflective of a robust positive association between trial-wise fixation count and similarity indices in younger but not older adults. The interaction in the LOC was not significant, exhibiting an age-invariant relationship between fixation count and LOC similarity indices. Thin lines illustrate random intercepts and slopes, bold lines illustrate the across-subject intercepts and slopes.

Turning to object trials, Figure 5C illustrates those object-selective parcels where fixation count covaried (positively or negatively) with similarity indices across the age groups, using the same formula described above. Negative associations were restricted to those regions that also exhibited below-zero object similarity (see illustrated in blue; Figure 4A) and are thus not discussed further. Positive across-age associations were present in the lateral occipital complex. An age group by fixation count interaction was evident in a single parcel within the posterior/inferior temporal gyrus (black outline in Figure 5C), which demonstrated a significant relationship in younger, but not older adults. Except for this one parcel, interaction effects were otherwise non-significant, indicating that trial-wise fixation count was positively associated with object selectivity in an age-invariant manner in most of the parcels that encompass the LOC.

Lastly, we performed analogous LME analyses at the ROI level. As noted in methods, the statistics from the ROI analyses reported below are derived from random intercept-only models to remain consistent with the whole-brain parcellation analyses. However, as we also noted, the inclusion of random slopes did not alter any of the statistical outcomes presented below (models employing random intercept and slopes are shown in Figure 5D for illustration purposes). Focusing first on the scene trials, we found that trial-wise fixation count significantly interacted with age group in predicting selectivity in the PPA (t = 2.966, p = 0.003), demonstrating that the strength of the association between PPA similarity and eye movements differed between the two age groups (Note that fixation count was a significant predictor of PPA selectivity even with the interaction included in the model (p < 0.001)). However, given that the relationship between selectivity and fixation count appeared to differ between the two age groups, we performed a follow-up analysis separately for younger and older adults (*PSA similarity ∼ fixation count + (1 | subject*). These two models revealed a significant positive relationship between PPA selectivity and fixation count in younger adults (t = 4.985, p < 0.001) but only a marginally significant relationship in the older adult group (t = 1.917, p = 0.055).

During object trials, the interaction between fixation count and age group was not significant (t = 1.307, p = 0.191), suggesting that the strength of the association between LOC similarity and eye movements did not differ between the two age groups. Therefore, across all participants, trial-wise fixation counts positively predicted LOC similarity (with interaction in the model: t = 2.537, p = 0.011; without interaction: t = 2.685, p = 0.007). In summary, the ROI analyses provided findings that were highly convergent with the whole brain analyses: the relationship between eye movements and PSA-based metrics of neural selectivity appears to be age-invariant in the LOC, but age-dependent in the PPA.

### Higher neural selectivity is associated with better subsequent memory for scenes in the PPA, but not for objects in the LOC

Two binomial generalized LMEs were employed to examine the relationship between subsequent memory success, age group, fixation count, and PSA similarity in the PPA (scene trials) and LOC (object trials). As noted in Methods, given that no significant interactions with age group were identified in any of the generalized LMEs reported below (ps > 0.163), and to avoid overfitting, the syntax used to generate all statistical outcomes was: *memory bin ∼ age group + PSA similarity + fixation count + (1 | subject)*. Memory success was binarized into correct (source correct) and incorrect trials (source incorrect, don’t know, item miss) prior to analysis. Significant predictors of subsequent memory success for scene trials were age group (z = 2.587, p = 0.010) and PPA similarity (z = 2.330, p = 0.020; higher similarity = higher probability of correct memory), but not fixation count (z = 0.765, p = 0.444). For the object trials, age group was a significant predictor of memory success (z = 4.435, p < 0.001), while LOC similarity and fixation count were not significant (z = 1.451, p = 0.147 and z = 0.797, p = 0.425). These results indicate that age group and scene-selectivity in the PPA explain unique sources of variance when predicting subsequent memory success for scene trials. On the other hand, object-selectivity in the LOC did not predict memory success above and beyond the variance associated with age group.

## Discussion

The present study examined possible associations between age differences in neural selectivity and eye movements. As reported previously (see Introduction), age effects on selectivity were most robust in scene-selective cortex. Importantly, the relationship between trial-wise estimates of neural selectivity and fixation counts was qualified by age group and stimulus category. While we identified an age-invariant positive association between selectivity and fixation counts during object viewing, the association was age-dependent during scene viewing, with a significant effect in young adults only. Below, we explore the significance of these findings for the understanding of the mechanisms that underpin age-related neural dedifferentiation and its impact on age differences in episodic memory performance.

Focusing first on behavioral performance, vividness ratings during encoding, as well as item recognition and source memory during retrieval, were lower in older than younger adults. These results align with the extensive prior literature documenting age-related declines in recollection, although we note that familiarity and item recognition are frequently reported to be spared in older age (for review, see Koen and Yonelinas, 2014). Given that vividness ratings at encoding have been reported to correlate positively with metrics of recollection derived from a subsequent memory test (Alghamdi & Rugg, 2020), at least some of the age-related variance in source memory performance may be attributable to the older adults’ tendency to form less vivid word-image associations at encoding (at least under the assumption that young and older adults had a common internal scale for vividness).

Age-related neural dedifferentiation has consistently been reported in scene-selective cortical regions, whereas dedifferentiation in object-, word-, and face-selective cortex is relatively elusive (for reviews, see Koen & Rugg, 2019; Koen et al., 2020). As we have argued previously (Srokova et al., 2020, 2024), scene-selective cortical areas, such as the PPA, are strongly modulated by perceptual complexity and attentional factors (e.g. Aminoff et al., 2013). This raises the possibility that lower selectivity for scene stimuli may be a consequence of a decline in the ability to process and bind the multiple spatially distributed elements that comprise the stimulus event. Alternatively, scene-related neural dedifferentiation could be a consequence of diminished availability of domain-general attentional resources and weakened attentional control of scene processing in older adults (c.f., Serences et al., 2004; Gazzaley et al., 2005; Bouhassoun et al., 2022).

The finding that older adults make more gaze fixations during static image viewing has been reported previously (Firestone et al., 2007; Heisz and Ryan, 2011; Liu et al., 2018; Wynn et al., 2021; Mazloum-Farzaghi et al., 2023). Indeed, it has been suggested that eye movement patterns in healthy older adults are similar, although less severe, to those observed in amnesia and age-related neurodegenerative disease. This observation prompted the hypothesis that differences in eye movement patterns are functionally related to memory deficits and hippocampal dysfunction (for reviews, see Ryan et al., 2020; Ryan & Shen, 2020; Wynn et al., 2020). Indeed, it is well-documented that exploratory eye movements are associated with hippocampal activity (e.g., Ringo et al., 1994; Jutras et al., 2013; Kragel et al., 2020; also see Meister & Buffalo, 2016). This has reinforced the idea that visual exploration and episodic memory operate within a dynamic feedback loop in which eye movements represent a critical component of hippocampally-mediated memory formation. Specifically, eye movements directly determine what becomes encoded into memory, and hippocampally-mediated mnemonic signals arising from previously viewed stimuli guide fixation sequences to allow subsequently viewed visual features to be bound into a coherent memory representation of the visual scene (Hannula et al., 2010; Voss et al., 2017).

In the present study, whole-brain parametric modulation analyses confirmed that, across age groups, fixation counts positively covaried with fMRI BOLD signals across the canonical scene-and object-selective cortical regions. This is consistent with prior studies indicating that eye movements modulate fMRI BOLD signals in early visual, parahippocampal, and occipito-temporal cortex (e.g., Henderson & Choi, 2015; Henderson et al., 2020; Liu et al., 2017, 2018; 2020; Ladyka-Wojcik et al., 2022). Of importance, we found that trial-wise fixation counts covaried with trial-wise estimates of scene- and object-selectivity. Crucially, however, while this relationship was age-invariant for objects in object-selective regions, the relationship in scene-selective cortex during scene viewing was evident in younger adults only.

Prior studies examining age differences in the relationship between simultaneously acquired fMRI BOLD signals and eye movements are sparse. However, an age-related weakening of this relationship during face viewing was reported by Liu and colleagues (2018). As in the present and prior behavioral studies, Liu and colleagues also reported an age-related elevation of fixation counts. Thus, the balance of the neuroimaging and behavioral evidence is consistent with the notion that more frequent fixations reflect a compensatory strategy to mitigate potential deficits in extracting visual information and binding it into memory (Olsen et al., 2016; Ryan et al, 2020). We emphasize that the relationship between neural selectivity and eye movements is unlikely to be unidirectional. Low fidelity of visual representations in higher-level visual cortex may be associated with degraded stimulus information that less efficiently guides sequences of gaze fixations during encoding. In return, inefficient sampling of the information might further degrade the stimulus representation and exacerbate age-related decline in neural selectivity.

Exploratory analyses of pupil size revealed smaller baseline-corrected pupil response in older relative to younger adults during word and image viewing, consistent with evidence of a reduction in stimulus-evoked pupil dilation in older adults (Zhao et al., 2019). Pupil dilation is thought to reflect task-evoked increases in processing load and effortful attention (for review, see Joshi & Gold, 2020). Pupil size has been shown to covary with neural firing rate in the locus coeruleus (LC), a brainstem nucleus implicated in arousal and attention (Varazzani et al., 2015; Joshi et al., 2016). Therefore, it is possible that age differences in pupil dilation result from degeneration of noradrenergic neurons in the LC (Mather & Harley, 2016), perhaps resulting in older adults devoting less attentional resources during word and image viewing. An alternative possibility is that present findings are a reflection of senile miosis - age-related reduction in baseline pupil size resulting from atrophy of the dilator muscles of the iris. Senile miosis may limit the dynamic range of pupillary responses in aging, leading to an underestimation of the cognitive efforts exerted by older individuals (Guillon et al., 2016; Herrmann & Ryan, 2024). We note, however, that the baseline-corrected pupil reflex evoked by the image onset did not differ across age groups, suggesting that restricted dynamic range cannot not fully explain the observed age differences in pupil size.

Lastly, the present study replicates prior findings that neural selectivity at encoding positively covaries with subsequent memory performance irrespective of age, and that this relationship is specific to the scene-selective PPA (Koen et al., 2019; Srokova et al., 2020; 2024). Here, we extend this result by demonstrating that the association remains significant when controlling for fixation counts, and that fixation counts did not predict memory success above and beyond neural selectivity (and neither did fixation predicted memory when neural selectivity was removed from the model, p = 0.324). This latter finding is particularly intriguing given that fixations at encoding typically predict memory success (Olsen et al., 2016; Damiano & Walther, 2019), underscoring the significance of neural selectivity to episodic memory performance. Nonetheless, it remains unknown why associations between neural selectivity and memory performance should be unique to scene images and the PPA. We conjecture that, given the role of the parahippocampal cortex in the formation of item-context associations (c.f., Diana et al., 2012; 2013), PPA selectivity may act as a general marker of the functional integrity of this region.

An important limitation of the present study lies in its cross-sectional design. Consequently, the conclusions drawn here cannot be unambiguously attributed to the effects of aging, and may instead reflect the influence of confounding variables, such as cohort effects (Rugg, 2017). We also cannot exclude the possibility that the moderating effects of age on the relationship between eye movements and neural activity arises from other mediating variables, such as the impact of aging on cerebral vasculature and hemodynamics (Fesharaki et al., 2024), or physiological changes to the eye that may affect eye-tracking in older adults (Gibson, 2013). Lastly, although low visual acuity was corrected with MR-safe glasses, we cannot rule out a contribution of other, related, factors such as age-related deterioration in contrast sensitivity (Owsley, 2016).

In conclusion, the present study provides novel evidence for a link between age-related neural dedifferentiation and eye movements. We confirm that age-related neural dedifferentiation is limited to scene-selective cortical regions, but our analyses go beyond prior findings by identifying a breakdown in the relationship between gaze fixations and scene selectivity in older adults. Thus, lower selectivity in scene-selective cortex may, at least in part, result from age differences in how eye movements are employed to sample visual stimuli. These findings offer new insights into the potential mechanisms underlying age-related neural dedifferentiation, thus shedding light on a putative determinant of age-related episodic memory decline.

## Acknowledgements

This research was supported by the National Institute of Aging Grants R56AG068149 and RF1AG039103, and BvB Dallas. S.S. is supported by the Postdoctoral training program of the Arizona Alzheimer’s Consortium, grant T32AG044402. The authors gratefully acknowledge Joshua Olivier, Ayse Aktas, and Eduardo Hernandez for their assistance with recruitment and neuropsychological assessments.

## Author contributions

S.S. and M.D.R. designed research, S.S. and N.S.S performed research, S.S. analyzed data, S.S. and M.D.R. wrote the paper.

## Conflicts of Interest

None

## Notes

### Competing Interest Statement

The authors have declared no competing interest.

## References

Alghamdi, S. A., & Rugg, M. D. (2020). The effect of age on recollection is not moderated by differential estimation methods. Memory, 28(8), 1067–1077.

Aminoff, E. M., Kveraga, K., & Bar, M. (2013). The role of the parahippocampal cortex in cognition. Trends in cognitive sciences, 17(8), 379–390.

Bailey, I. L., & Lovie-Kitchin, J. E. (2013). Visual acuity testing. From the laboratory to the clinic. Vision research, 90, 2–9.

Bouhassoun, S., Poirel, N., Hamlin, N., & Doucet, G. E. (2022). The forest, the trees, and the leaves across adulthood: Age-related changes on a visual search task containing three-level hierarchical stimuli. Attention, Perception, & Psychophysics, 1-12.

Carp, J., Park, J., Polk, T. A., & Park, D. C. (2011). Age differences in neural distinctiveness revealed by multi-voxel pattern analysis. Neuroimage, 56(2), 736–743.

Cassady, K., Gagnon, H., Freiburger, E., Lalwani, P., Simmonite, M., Park, D. C., … & Polk, T. A. (2020). Network segregation varies with neural distinctiveness in sensorimotor cortex. NeuroImage, 212, 116663.

Cassady, K., Gagnon, H., Lalwani, P., Simmonite, M., Foerster, B., Park, D., … & Polk, T. A. (2019). Sensorimotor network segregation declines with age and is linked to GABA and to sensorimotor performance. Neuroimage, 186, 234–244.

Chamberlain, J. D., Gagnon, H., Lalwani, P., Cassady, K. E., Simmonite, M., Seidler, R. D., … & Polk, T. A. (2021). GABA levels in ventral visual cortex decline with age and are associated with neural distinctiveness. Neurobiology of aging, 102, 170–177.

Chee, M. W., Goh, J. O., Venkatraman, V., Tan, J. C., Gutchess, A., Sutton, B., … & Park, D. (2006). Age-related changes in object processing and contextual binding revealed using fMR adaptation. Journal of cognitive neuroscience, 18(4), 495–507.

Damiano, C., & Walther, D. B. (2019). Distinct roles of eye movements during memory encoding and retrieval. Cognition, 184, 119–129.

de Chastelaine, M., Mattson, J. T., Wang, T. H., Donley, B. E., & Rugg, M. D. (2016). The relationships between age, associative memory performance, and the neural correlates of successful associative memory encoding. Neurobiology of aging, 42, 163–176.

Delis D.C., Kramer J.H., Kaplan E., Ober B.A., (2000) California verbal learning test, Ed. 2. San Antonio: The Psychological Corporation

Diana, R. A., Yonelinas, A. P., & Ranganath, C. (2012). Adaptation to cognitive context and item information in the medial temporal lobes. Neuropsychologia, 50(13), 3062–3069.

Diana, R. A., Yonelinas, A. P., & Ranganath, C. (2013). Parahippocampal cortex activation during context reinstatement predicts item recollection. Journal of Experimental Psychology: General, 142(4), 1287.

Ferris III, Frederick L., et al. “New visual acuity charts for clinical research.” American journal of ophthalmology 94.1 (1982): 91–96.

Fesharaki, N. J., Taylor, A., Mosby, K., Li, R., Kim, J. H., & Ress, D. (2024). Global Impact of Aging on the Hemodynamic Response Function in the Gray Matter of Human Cerebral Cortex. Human Brain Mapping, 45(18), e70100.

Firestone, A., Turk-Browne, N. B., & Ryan, J. D. (2007). Age-related deficits in face recognition are related to underlying changes in scanning behavior. Aging, Neuropsychology, and Cognition, 14(6), 594–607.

Gazzaley, A., Cooney, J. W., McEvoy, K., Knight, R. T., & D’esposito, M. (2005). Top-down enhancement and suppression of the magnitude and speed of neural activity. Journal of cognitive neuroscience, 17(3), 507–517.

Gipson, I. K. (2013). Age-related changes and diseases of the ocular surface and cornea. Investigative ophthalmology & visual science, 54(14), ORSF48–ORSF53.

Guillon, M., Dumbleton, K., Theodoratos, P., Gobbe, M., Wooley, C. B., & Moody, K. (2016). The effects of age, refractive status, and luminance on pupil size. Optometry and vision science, 93(9), 1093–1100.

Hannula, D. E., & Ranganath, C. (2009). The eyes have it: hippocampal activity predicts expression of memory in eye movements. Neuron, 63(5), 592–599.

Hannula, D. E., Althoff, R. R., Warren, D. E., Riggs, L., Cohen, N. J., & Ryan, J. D. (2010). Worth a glance: using eye movements to investigate the cognitive neuroscience of memory. Frontiers in human neuroscience, 4, 166.

Heisz, J. J., & Ryan, J. D. (2011). The effects of prior exposure on face processing in younger and older adults. Frontiers in Aging Neuroscience, 3, 15.

Henderson, J. M., Williams, C. C., & Falk, R. J. (2005). Eye movements are functional during face learning. Memory & cognition, 33(1), 98–106.

Henderson, J. M., & Choi, W. (2015). Neural correlates of fixation duration during real-world scene viewing: evidence from fixation-related (FIRE) fMRI. Journal of cognitive neuroscience, 27(6), 1137–1145.

Henderson, J. M., Goold, J. E., Choi, W., & Hayes, T. R. (2020). Neural correlates of fixated low-and high-level scene properties during active scene viewing. Journal of Cognitive Neuroscience, 32(10), 2013–2023.

Joshi, S., Li, Y., Kalwani, R. M., & Gold, J. I. (2016). Relationships between pupil diameter and neuronal activity in the locus coeruleus, colliculi, and cingulate cortex. Neuron, 89(1), 221–234.

Joshi, S., & Gold, J. I. (2020). Pupil size as a window on neural substrates of cognition. Trends in cognitive sciences, 24(6), 466–480.

Jutras, M. J., Fries, P., & Buffalo, E. A. (2013). Oscillatory activity in the monkey hippocampus during visual exploration and memory formation. Proceedings of the National Academy of Sciences, 110(32), 13144–13149.

Koen, J. D., & Yonelinas, A. P. (2014). The effects of healthy aging, amnestic mild cognitive impairment, and Alzheimer’s disease on recollection and familiarity: A meta-analytic review. Neuropsychology review, 24, 332–354.

Koen, J. D., & Rugg, M. D. (2019). Neural dedifferentiation in the aging brain. Trends in cognitive sciences, 23(7), 547–559.

Koen, J. D., Srokova, S., & Rugg, M. D. (2020). Age-related neural dedifferentiation and cognition. Current Opinion in Behavioral Sciences, 32, 7–14.

Kovach, (2007), https://github.com/UT-VisionMotionDecision/edfmex/blob/master/edfmex.m

Kragel, J. E., VanHaerents, S., Templer, J. W., Schuele, S., Rosenow, J. M., Nilakantan, A. S., & Bridge, D. J. (2020). Hippocampal theta coordinates memory processing during visual exploration. Elife, 9, e52108.

Kret, M. E., & Sjak-Shie, E. E. (2019). Preprocessing pupil size data: Guidelines and code. Behavior research methods, 51, 1336–1342.

Kriegeskorte, N., Mur, M., & Bandettini, P. A. (2008). Representational similarity analysis-connecting the branches of systems neuroscience. Frontiers in systems neuroscience, 2, 249.

Ladyka-Wojcik, N., Liu, Z. X., & Ryan, J. D. (2022). Unrestricted eye movements strengthen effective connectivity from hippocampal to oculomotor regions during scene construction. NeuroImage, 260, 119497.

Lalwani, P., Gagnon, H., Cassady, K., Simmonite, M., Peltier, S., Seidler, R. D., … & Polk, T. A. (2019). Neural distinctiveness declines with age in auditory cortex and is associated with auditory GABA levels. Neuroimage, 201, 116033.

Li, S. C., Lindenberger, U., & Sikström, S. (2001). Aging cognition: from neuromodulation to representation. Trends in cognitive sciences, 5(11), 479–486.

Liu, Z. X., Shen, K., Olsen, R. K., & Ryan, J. D. (2017). Visual sampling predicts hippocampal activity. Journal of Neuroscience, 37(3), 599–609.

Liu, Z. X., Shen, K., Olsen, R. K., & Ryan, J. D. (2018). Age-related changes in the relationship between visual exploration and hippocampal activity. Neuropsychologia, 119, 81–91.

Liu, Z. X., Rosenbaum, R. S., & Ryan, J. D. (2020). Restricting visual exploration directly impedes neural activity, functional connectivity, and memory. Cerebral cortex communications, 1(1), tgaa054.

Loftus, G. R. (1972). Eye fixations and recognition memory for pictures. Cognitive psychology, 3(4), 525–551.

Maass, A., Berron, D., Harrison, T. M., Adams, J. N., La Joie, R., Baker, S., … & Jagust, W. J. (2019). Alzheimer’s pathology targets distinct memory networks in the ageing brain. brain, 142(8), 2492–2509.

Mazloum-Farzaghi, N., Shing, N., Mendoza, L., Barense, M. D., Ryan, J. D., & Olsen, R. K. (2023). The impact of aging and repetition on eye movements and recognition memory. Aging, Neuropsychology, and Cognition, 30(3), 402–428.

Mather, M., & Harley, C. W. (2016). The locus coeruleus: essential for maintaining cognitive function and the aging brain. Trends in cognitive sciences, 20(3), 214–226.

Meister, M. L., & Buffalo, E. A. (2016). Getting directions from the hippocampus: The neural connection between looking and memory. Neurobiology of learning and memory, 134, 135–144.

Mumford, J. A., Davis, T., & Poldrack, R. A. (2014). The impact of study design on pattern estimation for single-trial multivariate pattern analysis. Neuroimage, 103, 130–138.

Olsen, R. K., Sebanayagam, V., Lee, Y., Moscovitch, M., Grady, C. L., Rosenbaum, R. S., & Ryan, J. D. (2016). The relationship between eye movements and subsequent recognition: Evidence from individual differences and amnesia. Cortex, 85, 182–193.

Owsley, C. (2016). Vision and aging. Annual review of vision science, 2(1), 255–271.

Park, D. C., Polk, T. A., Park, R., Minear, M., Savage, A., & Smith, M. R. (2004). Aging reduces neural specialization in ventral visual cortex. Proceedings of the National Academy of Sciences, 101(35), 13091–13095.

Park, J., Carp, J., Hebrank, A., Park, D. C., & Polk, T. A. (2010). Neural specificity predicts fluid processing ability in older adults. Journal of Neuroscience, 30(27), 9253–9259.

Park, J., Carp, J., Kennedy, K. M., Rodrigue, K. M., Bischof, G. N., Huang, C. M., … & Park, D. C. (2012). Neural broadening or neural attenuation? Investigating age-related dedifferentiation in the face network in a large lifespan sample. Journal of Neuroscience, 32(6), 2154–2158.

Payer, D., Marshuetz, C., Sutton, B., Hebrank, A., Welsh, R. C., & Park, D. C. (2006). Decreased neural specialization in old adults on a working memory task. Neuroreport, 17(5), 487–491.

Peirce, J., Gray, J. R., Simpson, S., MacAskill, M., Höchenberger, R., Sogo, H., … & Lindeløv, J. K. (2019). PsychoPy2: Experiments in behavior made easy. Behavior research methods, 51, 195–203.

Peitek, N., Siegmund, J., Parnin, C., Apel, S., Hofmeister, J. C., & Brechmann, A. (2018, October). Simultaneous measurement of program comprehension with fmri and eye tracking: A case study. In Proceedings of the 12th ACM/IEEE international symposium on empirical software engineering and measurement (pp. 1–10).

PsychoPy2: Experiments in behavior made easy. Behavior research methods, 51, 195–203.

R Core Team (2020) R: a language and environment for statistical computing. Vienna: R Foundation.

Raven, J., Raven, J.C., Courth, J.H. (2000) The advanced progressive matrices. In: Manual for Raven’s progressive matrices and vocabulary scales, Section 4. San Antonio: Harcourt Assessment.

Reitan, R.M., Wolfson, D. (1985) The Halstead-Reitan neuropsychological test battery: therapy and clinical interpretation. Tucson: Neuropsychological.

Ringo, J. L., Sobotka, S., Diltz, M. D., & Bunce, C. M. (1994). Eye movements modulate activity in hippocampal, parahippocampal, and inferotemporal neurons. Journal of Neurophysiology, 71(3), 1285–1288.

Rissman, J., Gazzaley, A., & D’Esposito, M. (2004). Measuring functional connectivity during distinct stages of a cognitive task. Neuroimage, 23(2), 752–763.

Rugg MD (2017). Interpreting age-related differences in memory-related neural activity. In Cognitive neuroscience of aging: Linking cognitive and cerebral aging, 2nd ed (pp. 183–203). Oxford University Press.

Ryan, J. D., & Shen, K. (2020). The eyes are a window into memory. Current Opinion in Behavioral Sciences, 32, 1–6.

Ryan, J. D., Shen, K., & Liu, Z. X. (2020). The intersection between the oculomotor and hippocampal memory systems: empirical developments and clinical implications. Annals of the New York Academy of Sciences, 1464(1), 115–141.

Sekuler, B., Bennett, P. J., Mamelak, M. A.(2000). Effects of aging on the useful field of view. Experimental aging research, 26(2), 103–120.

Serences, J. T., Schwarzbach, J., Courtney, S. M., Golay, X., & Yantis, S. (2004). Control of object-based attention in human cortex. Cerebral cortex, 14(12), 1346–1357.

Schaefer, A., Kong, R., Gordon, E. M., Laumann, T. O., Zuo, X. N., Holmes, A. J., … & Yeo, B. T. (2018). Local-global parcellation of the human cerebral cortex from intrinsic functional connectivity MRI. Cerebral cortex, 28(9), 3095–3114.

Sheng, J., Trelle, A. N., Romero, A., Park, J., Tran, T. T., Sha, S., … & Wagner, A. D. (2024). Top-down attention and Alzheimer’s pathology impact cortical selectivity during learning, influencing episodic memory in older adults. bioRxiv, 2024-12. 10.1101/2024.12.04.626911

Smith, A. (1982) Symbol digit modalities test (SDMT) manual. Los Angeles: Western Psychological Services.

Spreen O., Benton A.L. (1977) Neurosensory center comprehensive examination for aphasia. Victoria: Neuropsychology Laboratory.

Srokova, S., Hill, P. F., Koen, J. D., King, D. R., & Rugg, M. D. (2020). Neural differentiation is moderated by age in scene-selective, but not face-selective, cortical regions. ENeuro, 7(3).

Srokova, S., Aktas, A. N., Koen, J. D., & Rugg, M. D. (2024). Dissociative effects of age on neural differentiation at the category and item levels. Journal of Neuroscience, 44(4).

Varazzani, C., San-Galli, A., Gilardeau, S., & Bouret, S. (2015). Noradrenaline and dopamine neurons in the reward/effort trade-off: a direct electrophysiological comparison in behaving monkeys. Journal of Neuroscience, 35(20), 7866–7877.

Voss, M. W., Erickson, K. I., Chaddock, L., Prakash, R. S., Colcombe, S. J., Morris, K. S., … & Kramer, A. F. (2008). Dedifferentiation in the visual cortex: an fMRI investigation of individual differences in older adults. Brain research, 1244, 121–131.

Voss, J. L., Bridge, D. J., Cohen, N. J., & Walker, J. A. (2017). A closer look at the hippocampus and memory. Trends in cognitive sciences, 21(8), 577–588.

Wechsler, D. (1981) WAIS-R: Wechsler adult intelligence scale-revised. New York: The Psychological Corporation.

Wechsler, D. (2009) Wechsler memory scale, Ed 4. San Antonio: The Psychological Corporation.

Wechsler, D. (2011) The Test of Premorbid Function (TOPF). The Psychological Corporation, San Antonio, TX

Wynn, J. S., Buchsbaum, B. R., & Ryan, J. D. (2021). Encoding and retrieval eye movements mediate age differences in pattern completion. Cognition, 214, 104746.

Zebrowitz, L., Ward, N., Boshyan, J., Gutchess, A., & Hadjikhani, N. (2016). Dedifferentiated face processing in older adults is linked to lower resting state metabolic activity in fusiform face area. Brain research, 1644, 22–31.

Zhao, S., Bury, G., Milne, A., & Chait, M. (2019). Pupillometry as an objective measure of sustained attention in young and older listeners. Trends in hearing, 23, 2331216519887815.

Zheng, L., Gao, Z., Xiao, X., Ye, Z., Chen, C., & Xue, G. (2018). Reduced fidelity of neural representation underlies episodic memory decline in normal aging. Cerebral Cortex, 28(7), 2283–2296.

